# Atlas of telomeric repeat diversity in *Arabidopsis thaliana*

**DOI:** 10.1101/2023.12.18.572118

**Authors:** Yueqi Tao, Wenfei Xian, Fernando Rabanal, Andrea Movilli, Christa Lanz, Gautam Shirsekar, Detlef Weigel

**Affiliations:** Department of Molecular Biology, Max Planck Institute for Biology Tübingen, Tübingen 72076, Germany

## Abstract

Telomeric repeat arrays at the ends of chromosomes are highly dynamic but their repetitive nature and technological limitations have made it difficult to assess the variation in genome diversity surveys. Here we present a comprehensive characterization of the sequence variation immediately adjacent to the canonical telomeric repeat arrays at the very ends of chromosomes in 49 genetically diverse *Arabidopsis thaliana* accessions. We reveal several types of distinct telomeric repeat units and identify evolutionary processes such as local homogenization and higher-order repeat formation that shape diversity of chromosome ends. The identification of segmental duplications and at least one recombination event suggests a plausible history of telomerase-independent maintenance generation. By comparing largely isogenic samples, we are able to determine variant telomeric repeat number variation at both the germline and somatic levels. Analysis of haplotype structure uncovers chromosome end-specific as well as genetic group-specific patterns in telomeric repeat diversity and provides evidence for linkage disequilibrium between repeat arrays and their adjacent non-coding regions. Together, our findings illustrate the fine-scale telomeric repeat spectrum in *A. thaliana*, expanding our knowledge of the evolution of chromosome ends.

## INTRODUCTION

Telomeric repeat arrays are found at the termini of most eukaryotic chromosomes (Churikov and Price 2008). The very ends of the arrays, known as telomeres (Chan and Blackburn 2004), commonly consist of canonical units with the formula (T)_x_(A)_y_(G)_z_ and act as functional caps that protect chromosome ends from degradation and fusion (Fulnecková et al. 2013; Verdun and Karlseder 2007). These canonical repeats are being synthesized from an RNA template by telomerase, which ensures their sequence conservation (Schrumpfová and Fajkus 2020). In contrast to these highly conserved repeats, the immediately following sequences often include degenerate and variant telomeric repeats (Wallberg et al. 2019; Richards et al. 1992; Allshire et al. 1989), which differ from the canonical unit in one or more base substitutions or small insertions and deletions (indels; Lee et al. 2014). The composition of the variant repeats displays remarkable heterogeneity within the same genetic group and among different chromosome ends (Tham et al. 2023; Mizuno et al. 2008; Baird et al. 2000), raising questions as to the evolutionary mechanisms that generate and maintain this diversity (Mendez-Bermudez et al. 2009; Pickett et al. 2004). This telomere-adjacent region serves as a transition zone between the telomere and the rest of the chromosome that contains genes and other genetic elements (Churikov and Price 2008). Specific types of variant telomeric repeats have been implicated in determining methylation state (Farrell et al. 2022), protein binding (Wang et al. 2023) and formation of G-quadruplexes (Lee and Kim 2009). A comprehensive understanding of the evolutionary dynamics and functional significance of telomeres and telomere-adjacent regions must therefore begin with thorough knowledge of variation in the composition of telomeric repeats.

*Arabidopsis thaliana* has a seven base pair canonical unit TTTAGGG, that is the dominant telomeric unit in many other plant species as well (Peska and Garcia 2020; Richards and Ausubel 1988). The presence of variant telomeric repeats in *A. thaliana* was first established with a yeast artificial chromosome strategy (Richards et al. 1992). Subsequently, sequencing of PCR products revealed the heterogeneity of variant repeats from individual chromosome ends (Wang et al. 2010; Kuo 2006). Variant repeats have also been directly observed in unassembled sequencing reads (Choi et al. 2021), and they have been identified by partially assembling four chromosome ends in the Col-0 accession from Illumina short reads (Farrell et al. 2022). However, the highly repetitive nature of telomeric regions, the presence of identical sequences shared between repeat-adjacent regions, as well as large interstitial telomeric arrays in other parts of *A. thaliana* genomes create ambiguity when mapping reads that are only hundreds base pairs long to specific positions of the genome (Olson et al. 2023; Teano et al. 2023; Heacock et al. 2004). As a result, variation in telomeric repeat content at *A. thaliana* chromosome ends remains largely uncharacterized and has been ignored in diversity studies.

New single-molecule sequencing methods, generating reads of more than 10 kilobases (kb) in length, which exceeds the size of full-length telomeric repeat tracts and extend into unique repeat-adjacent regions, can overcome the challenges of reconstructing full telomeric sequences (Grigorev et al. 2021). However, although several *A. thaliana* genome assemblies have now been published (Hou et al. 2022; Wang et al. 2022; Naish et al. 2021), they have largely ignored the telomeric sequences apart from confirming that telomeres are structurally present at most chromosome ends. Pacific Biosciences High Fidelity (PacBio HiFi) sequencing is particularly well suited for reliable base calling in low-complexity telomeric repeats (Tan et al. 2022). In addition, the circular sequencing mode of HiFi sequencing, wherein each DNA molecule is sequenced multiple times, allows us to confidently characterize somatic information such as repeat number variation in the telomeric regions, which is obscured in assemblies (Wenger et al. 2019; Loomis et al. 2013).

In this study, we provide a high-resolution description of telomeric repeats for all ten chromosome ends in *A. thaliana*. We identify numerous types of variant repeats and previously undescribed sequence structures, some of which may reveal an uncharacterized evolutionary history of telomerase-independent maintenance. We also investigate telomeric repeat number variation at germline and somatic levels. We illustrate chromosome end-specific and genetic group-specific patterns of repeat haplotypes along with linkage disequilibrium between telomeric repeat arrays and their adjacent non-coding regions. Our findings significantly expand the collection of repeats derived from canonical telomeric repeats and telomeric sequence features in *A. thaliana*, setting the stage for a deeper understanding of the evolutionary mechanisms acting on them.

## RESULTS

### Profiling telomeric regions in *A. thaliana*

To investigate the sequence content of telomeric regions, defined here as canonical telomeric repeats, adjacent variant and degenerate telomeric repeats as well as any unique sequences interspersed in these repeats, HiFi reads from 49 *A. thaliana* accessions of diverse geographic origin were used (Supplemental Fig. S1). Data from 46 accessions were publicly available, and three additional accessions were sequenced for this study (Supplemental Table S1). For each accession, HiFi reads were extracted for each chromosome end, and the composition of repeat units was determined for each read (Supplemental Fig. S2A). For the ends of the p-arms of chromosome 2 and 4 (hereafter, chr2p and chr4p), which remain incompletely assembled due to large 45S ribosomal DNA (rDNA) tandem arrays that are immediately adjacent to the telomeres (Copenhaver and Pikaard 1996), reads could be assigned to two groups but could not be unambiguously assigned to chr2p or chr4ps (Supplemental Fig. S2B).

Starting from the centromere-proximal side, the telomeric regions typically start with a stretch of degenerate repeats, followed by variant repeats and finally canonical repeats, all of which were in the same head-to-tail arrangement (Fig. 1A). The most obvious exceptions to this general pattern were chr2p and chr4p ends, where only five accessions had variant repeats. Additionally, 19 accessions contained non-repeat fragments within the repeat arrays, and these are described in detail below.

**Figure 1.**
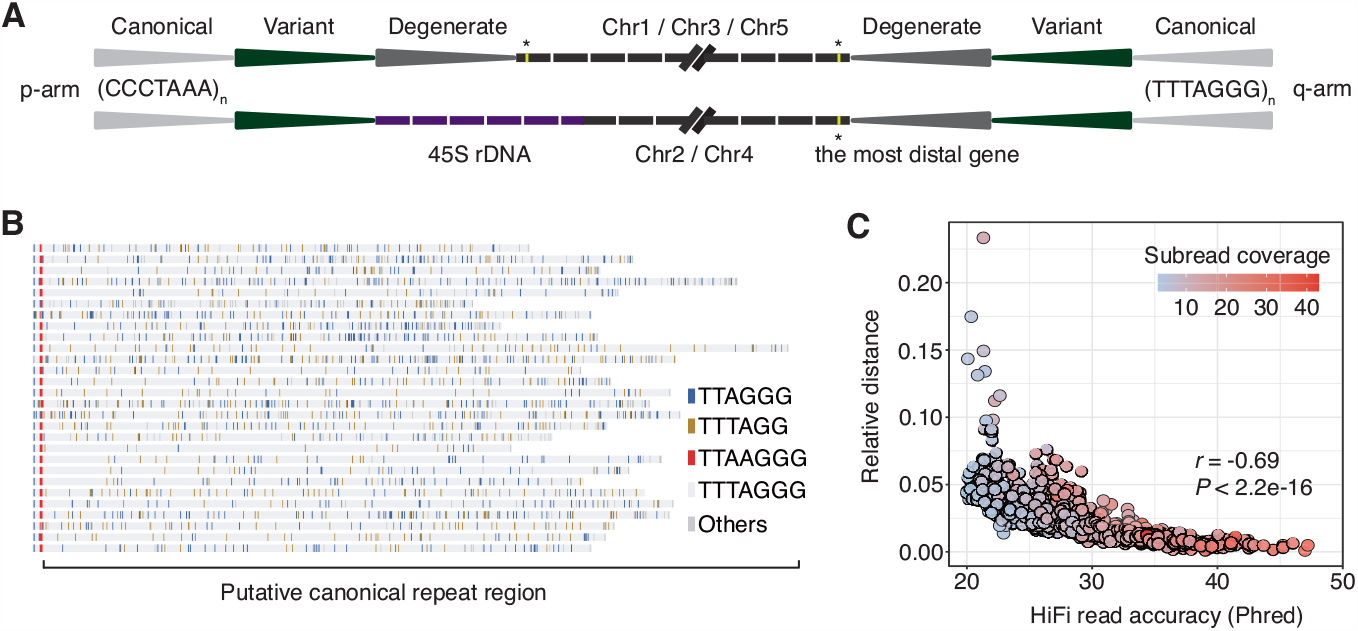
Overview of the telomeric repeat regions in *A. thaliana*. **(A)** Schematic representation of the different types of telomeric repeats at non-rDNA and rDNA chromosome ends. **(B)** Alignment of HiFi reads showing the entire telomeric repeat array in chr3q of NA2212 accession from degenerate, variant to canonical repeats (from left to right). **(C)** Correlation between relative edit distance from expected canonical repeat sequence and entire read accuracy.

Alignments of the HiFi reads from individual chromosome ends revealed that every read harbored many 1-bp indels, primarily occurring in the putative canonical repeats towards the ends of the reads, usually replacing TTTAGGG with either TTAGGG or TTTAGG (Fig. 1B; Supplemental Fig. S3). By analyzing the correlation between HiFi read accuracy, the number of full-pass subreads, and the proportion of indels for each read, a statistically significant negative correlation was found between the proportion of indels and both read accuracy (*P* < 2.2e-16, Pearson’s *r* = -0.69; Fig. 1C) and subread coverage (*P* < 2.2e-16, Pearson’s *r* = -0.42). This result suggests that the occurrence of indels is influenced by the read quality. Different from a previous study that interpreted indels supported by a single read as genuine variants (Grigorev et al. 2021), we consider these indels to be short homopolymer run errors (TTT>TT or GGG>GG), a known issue with HiFi reads (Lal et al. 2021; Loomis et al. 2013). Therefore, the region beginning at the last conserved variant repeat until the read end was defined as the canonical TTTAGGG repeat region. Because it was deemed to be devoid of consistent variation, this region was not further considered in the remainder of analyses.

### Hypervariable composition of telomeric repeat arrays

Using the extracted reads, we generated consensus sequences of degenerate and variant telomeric repeats for each chromosome end in the 49 accessions. To obtain a first overview of variation, the 15 most enriched repeat types were visualized (Fig. 2). The length of the arrays varied from zero to 3,384 bp (chr1p of Ey15-2). Of the 392 non-rDNA chromosome ends, 371 had variant repeat arrays, with lengths from 6 to 3,384 bp. Of the 98 rDNA ends, only 5 had variant repeat arrays, with lengths from 52 to 658 bp. A total of 385 distinct repeat units, ranging in size from 2 to 15 bp, were identified (Supplemental Table S2). Of these, 126 units (32.7%) occurred only once. The canonical repeat, which was interspersed among arrays of variant repeats, had the highest frequency with 13.3%. A few chromosome ends, including chr5p of Cas-0 as the most extreme example, had a large number of consecutive TTCAGGG repeats, which has been proposed to result from a mechanism known as alternative lengthening of telomeres (ALT; Marzec et al. 2015). It should be noted that the count of distinct repeat types greatly relies on our definition of a unit. For example, the sequence TTTAGGATTAGGG could be considered as being composed of two variant repeats, TTTAGGA and TTAGGG, or TTTAGG and ATTAGGG. Therefore, we use the repeat types as a set of markers for studying the overall organization of telomeric sequences and believe that there is no need to excessively focus on the specific sequence content of individual units, especially rare ones.

**Figure 2.**
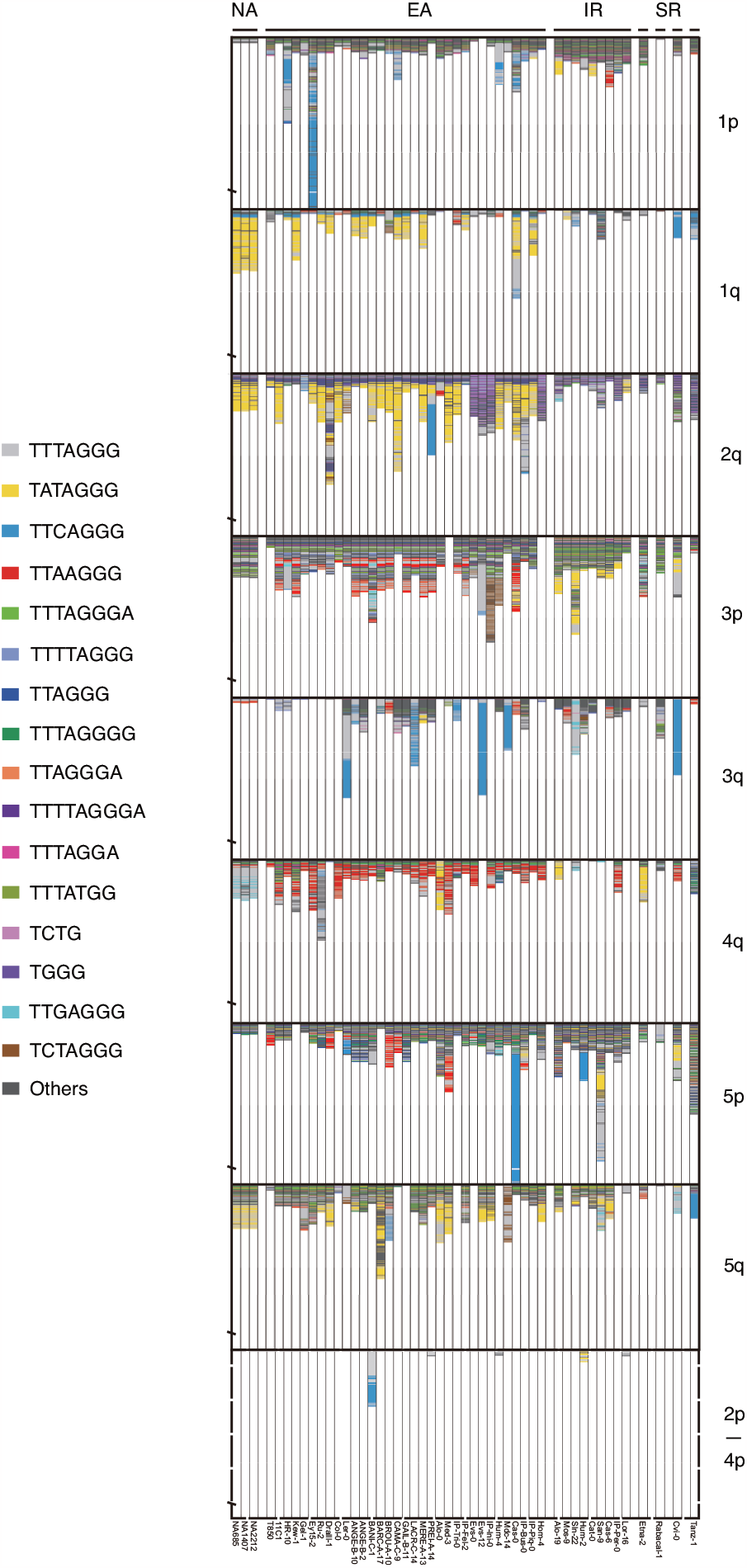
Degenerate and variant (from top down) telomeric repeat arrays at 10 chromosome ends across all 49 accessions. The top 15 most enriched units are highlighted by different colors. Bar height represents the length of the degenerate and variant repeat region. The slash denotes the location of the 300th repeat. Genetic groups and accessions are indicated at the top and at bottom of the graph, respectively (NA, North American; EA, Eurasian; IR, Iberian relict; SR, single relict).

As an aside, the template sequence of the telomerase RNA, 5′-CUAAACCCU-3′ (Song et al. 2019), encoded on chromosome 2, was identical in all 49 accessions (Supplemental Fig. S4).

Although the sequence content of telomeric regions was highly dynamic, there were five main patterns of sequence variation and most non-rDNA chromosome ends, 341 of 392, showed more than one of these patterns. The simplest pattern was represented by arrays in which different repeat types occurred only once, such as chr1q of Cat-0 (Fig. 2). The second pattern most likely resulted from monomer homogenization, such as chr1p of Alo-19, where a single unit, TATAGGG, was repeated consecutively 15 times (Supplemental Fig. S5A). The remaining patterns constituted higher-order repeats (HORs; Garrido-Ramos 2017). In the simplest case, such as chr3q of IP-Tri-0, two to four units made up a block that was then repeated multiple times (Supplemental Fig. S5B). A more elaborate pattern had multiple monomers (arbitrarily defined as ≥ 5 here) that were repeated several times. For example, in chr2q of IP-Per-0, five distinct units formed a block and were repeated five times, with all five blocks being identical (Supplemental Fig. S5C). The final pattern also featured HORs, but with mutations distinguishing the individual HORs. For instance, in chr2q of Cvi-0, the HOR array consisted of five units repeated eight times with five of these deviating from the consensus (Supplemental Fig. S5D).

When comparing pairs of accessions, the majority of sequence differences between specific chromosome ends fell into three major groups (Fig. 2). In the first category, sequences were highly similar to each other, as seen in chr5p of 11C1 and HR-10. In the second group, sequence composition was similar, but accessions were distinguished by the number of HORs, such as chr3p of IP-Tri-0 and IP-Fel-2. These two categories were mainly observed with pairs from the same genetic group such as Eurasian or Iberian relicts. The third group, sequence divergence, was observed not only in unrelated accessions but also in pairs from exactly the same local population, such as Evs-0 and Evs-12, where all eight non-rDNA chromosome ends were dissimilar.

### Non-repeat sequences in telomeric repeat arrays

Nineteen accessions had non-repeat sequences within the repeat array (Supplemental Table S3). Except for seven unclassified sequences ranging in length from 42 to 453 bp, the others could typically be divided into three different types. Firstly, organellar DNA or rDNA insertions. In chr1p, nine accessions had a 110-bp mitochondrial DNA insertion (Supplemental Fig. S6), which has been reported previously (Kuo et al. 2006), while chr2q of Cvi-0 contained a 102-bp chloroplast DNA insertion. A 5,088-bp 45S rDNA sequence was embedded in the telomeric tract in chr2q of Gel-1. In the second type, there were non-repeat fragments that were associated with repeat array duplications. In chr2q of Dralll-1, there was a HOR that included a 244-bp unique sequence, and the entire arrangement was repeated three times (Fig. 3A). The 244-bp fragment is identical in all three HOR copies, while the repeat array exhibits a few variants. Similarly, in chr3q of Cas-6, the HOR included a 3,865-bp unique sequence, and the entire sequence was repeated twice (Fig. 3B). The third type was exemplified by chr3q of Hum-2, where the repeat array was interrupted by a 495-bp non-repeat fragment, which was identical in sequence to a fragment adjacent to the array of variant telomeric repeats of chr5q in the same accession (Fig. 3C). The distal part of chr3q closely resembled the repeat array of chr5q, suggesting gene conversion or recombination. These large-scale variants of the typical telomeric repeat arrays have not been reported in *A. thaliana*. The only other similar example that we are aware of comes from *Caenorhabditis elegans* (Kim et al. 2019), and it was also discovered using PacBio single-molecule sequencing.

**Figure 3.**
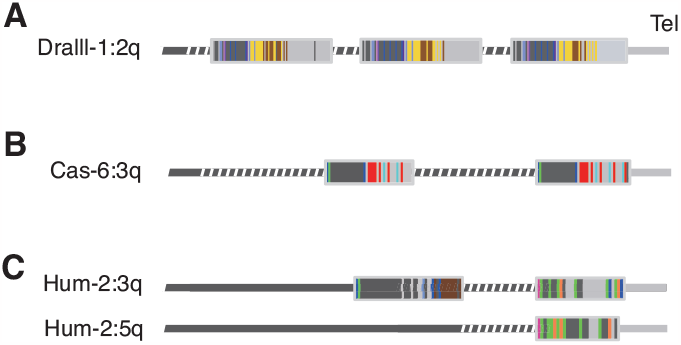
Representation of telomeric repeat arrays that include unique sequence fragments. Darker gray bars indicate repeat-adjacent regions. Dashed lines indicate unique sequences that interrupt arrays of variant repeats (indicated in color). The telomere (Tel) is on the right-hand side in all three cases (indicated in light gray). **(A)** Chr2q of Dralll-1. **(B)** Chr3q of Cas-6. **(C)** A 495-bp unique sequence in chr3q of Hum-2 is also found in chr5q.

### Repeat number variation between closely related individuals and in somatic tissues

To examine variability in the telomere regions in a more fine-grained manner, two collections of datasets from very closely related individuals were employed. The first collection came from the lineage of North American accessions known as haplogroup-1 (HPG1), which form a clade of natural mutation accumulation lines whose common ancestor lived about 400 years ago (Exposito-Alonso et al. 2018). In parallel, three independent sequencing datasets of the Col-0 accession that had been recently published were investigated (Rabanal et al. 2022; Wang et al. 2022; Naish et al. 2021). This also offered an opportunity to examine intra-dataset variation in more detail. We therefore report not merely the most common repeat array length, but present the full data for all HiFi reads.

Among the three HPG1 accessions, repeat number variation was found, but no major differences were observed in repeat type. Specifically, four of eight non-rDNA chromosome ends were significantly different in lengths of degenerate and variant repeat regions, with medians differing from 7 to 51 bp (Fig. 4A). There was also substantial variation in repeat number among the HiFi reads from a single accession. The greatest one, from 396 to 569 bp, corresponding to approximately 25 repeat units, was observed at chr4q of NA1407 (Fig. 4A).

**Figure 4.**
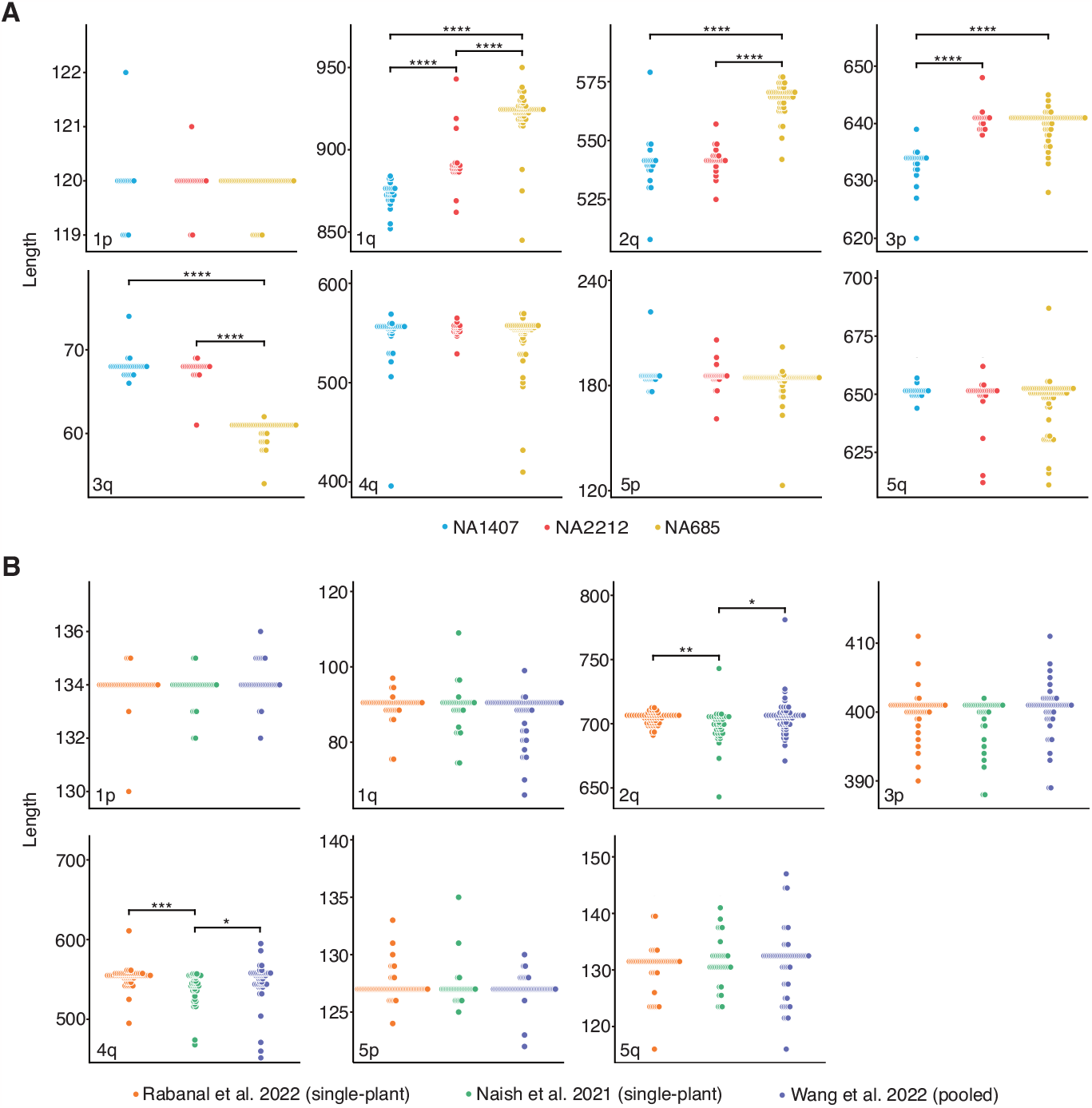
Plots showing variation in the length of degenerate and variant telomeric repeat regions in sets of three closely related samples. Dots represent individual HiFi reads. Statistically significant differences are indicated (****P<0.00001, ***P<0.0001, **P<0.001, *P<0.01). **(A)** Comparison of the three HPG1 accessions. **(B)** Comparison of the three Col-0 datasets.

In the Col-0 accession, the array of telomeric repeats of chr3q was found to exclusively consist of canonical repeats and it was therefore excluded from this analysis. For the remaining seven non-rDNA chromosome ends, there was no difference in variant types. Regarding repeat number variation, at two of seven chromosome ends, one dataset differed significantly in length distribution from the other two datasets, with median differences of 7 bp and 11 bp (Fig. 4B). These two chromosome ends had also the longest repeat arrays. For within-dataset length variation, chr4q was the one with the greatest difference between the shortest and longest arrays of degenerate and variant repeats, at 184 bp, roughly equivalent to 26 repeats (Fig. 4B). While four of seven chromosome ends differed significantly in the degrees of variability among the three Col-0 datasets (Supplemental Fig. S7), these differences were not attributable to the pooled-sequencing dataset. Thus, differences in sequencing strategy should not affect our conclusions regarding the 49 diverse datasets we used, which had been generated by a combination of pooled and single-plant sequencing.

### Haplotype structure of telomeric repeat arrays and the adjacent non-coding regions

To facilitate the comparison of haplotypes across the telomeric arrays, we implemented a repeat compression process to mitigate the impact of repeat number variation, which is likely to change more quickly than the overall arrangement and presence of variant repeats (Supplemental Fig. S8). The compressed sequences were used to perform a pairwise sequence similarity analysis based on the Levenshtein distance (L-distance; Levenshtein, 1966). The result confirmed the visual impression from Figure 2, that there is on average more similarity between the same chromosome end of different accessions than between different chromosome ends (Supplemental Fig. S9A; *P* < 2.2e-308, Wilcoxon test). The results also showed an overall lower L-distance within the same genetic group compared to between different genetic groups (Supplemental Fig. S9B; *P* < 6.01e-59, Wilcoxon test).

To examine whether these haplotype patterns extended beyond the telomeric repeat regions, we also looked at their adjacent non-coding regions. Non-coding sequences, which varied in length from zero to 16,542 bp, were defined as the sequence between the most distal gene and the last variant repeat of each chromosome end (Supplemental Table S4). Next, neighbor joining (NJ) clustering was conducted based on the multi-sequence alignment of these non-coding regions from each chromosome end. A merged matrix of repeat arrays and non-coding regions was generated, using the accession order from the NJ exercise, to reveal the correlation between the two (Fig. 5). Strong linkage between telomeric repeats and their adjacent non-coding regions were present at both coarse and fine resolution.

**Figure 5.**
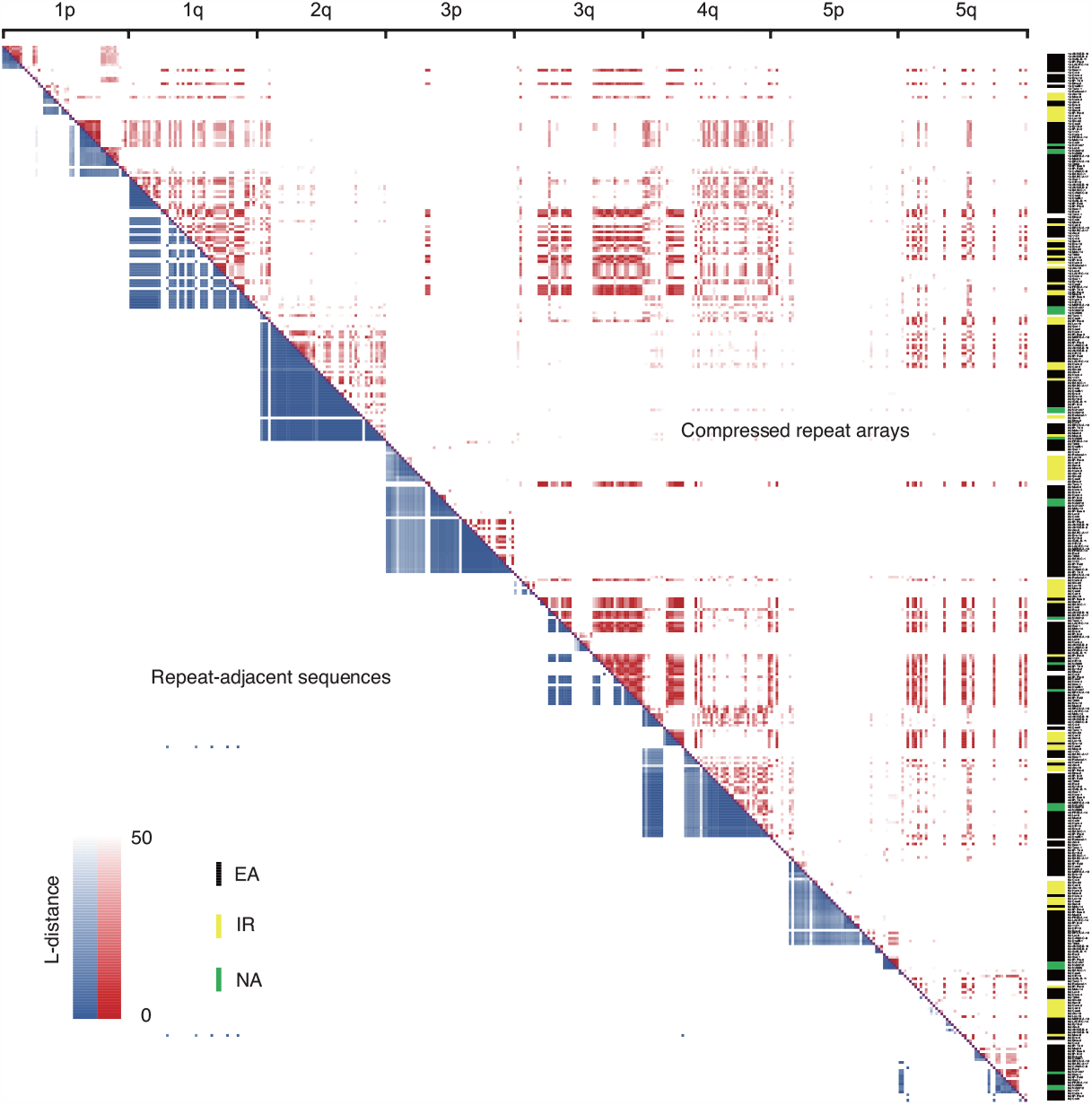
Heatmap of pairwise L-distances of the telomeric repeat arrays (upper triangle) and the repeat-adjacent sequences (lower triangle). Membership of accessions in different genetic groups is denoted by the colors on the right (EA, Eurasian; IR, Iberian relict; NA, North American).

Here we take chr1p, the chromosome end wherein the clearest pattern, as an example. Within chr1p, the non-coding sequences could be divided into three main haplotype clusters based on substantial sequence differences. Two of these clusters corresponded to a single repeat array haplotype each, while the third cluster was further subdivided by three repeat array haplotypes. Each subgroup of the third cluster, which corresponded to a specific type of repeat array, shared several single polymorphic sites in its non-coding sequences. Three distinct haplotype clusters were also apparent in chr3q, and two in chr4q. The remaining chromosome ends also showed linkage between repeat arrays and non-coding sequences, although this was less obvious in cases with longer repeat arrays and therefore higher absolute L-distance.

In addition to linkage disequilibrium, the matrix provided a direct support for our statistical results regarding the chromosome end-specific and genetic group-specific patterns (Fig. 5). Haplotypes from the same chromosome end clustered together. It is worth mentioning here that although some similarities in haplotype membership across chromosome ends were observed, they were mainly due to small sizes of repeat arrays only allowing for small edit distances. For all chromosome ends, Eurasian accessions were more likely to cluster with other Eurasian accessions, and the same was true for Iberian relict accessions. The three North American accessions formed their own cluster for some chromosome ends or clustered closely with Eurasian accessions for some other chromosome ends, which was not unexpected, given that the source population was in Europe (Exposito-Alonso et al. 2018).

## DISCUSSION

Our study provides a base-level view of the patterns of degenerate and variant telomeric repeats at the chromosome ends of 49 geographically diverse accessions of *A. thaliana*. We were able to unambiguously anchor these repeats to each chromosome end and greatly extend knowledge of telomere-adjacent sequence variation in this species (Kuo et al. 2006; Richards et al. 1992). Of note, our results overturned the previous conclusion that there are no variants at the two chromosome ends that cap the large 45S rDNA repeat arrays (Copenhaver and Pikaard 1996). The comprehensive picture presented here both leads to hypotheses for how sequence diversity at telomeres is generated and maintained, and it will help to explore what specific functions are associated with particular types of telomeric repeats (Dubocanin et al. 2022; Mendez-Bermudez et al. 2009).

There is ample evidence for local homogenization of telomeric repeats and formation of higher-order repeats (HORs), as well as repeat number variation in somatic cells and between closely related individuals, all typical characteristics of non-coding minisatellite regions (Garrido-Ramos 2017; Boán et al. 2004). The obvious scenario is that only the most distal portions of the canonical repeats, at the very ends of the chromosomes, are maintained by telomerase and remain thus uniform (Song et al. 2019). More centromere-proximal portions are maintained by conventional DNA replication and can sustain mutations, becoming first variant repeats and eventually degenerate repeats over time (Kuo et al. 2006). In this scenario, the variant and degenerate repeats are minisatellite units of about 7 bp, and the extensive patterns of apparent repeat expansion and contraction can be explained by replication slippage and unequal crossing over (Symonds and Lloyd 2003). Two other forces shaping variant repeats have been considered in previous studies, and our analyses cannot rule out that they play a minor role as well. Variant repeats could in principle be caused by variation in the RNA template. While we detected no sequence differences at the previously reported locus for the canonical RNA template (Fajkus et al. 2019; Song et al. 2019), we cannot exclude the existence of other loci that contribute a minor amount of alternative templates (Závodník et al. 2023). Alternatively, variants could arise during reverse transcription (Gout et al. 2013), introducing variants into the newly added repeats at the most distal end of the array. Such errors will cause telomere elongation by telomerase to fail, with alternative mechanisms for telomere maintenance eventually taking over.

Our haplotype analysis revealed both chromosome end-specific and genetic group-specific patterns of degenerate and variant telomeric repeat arrays. Accessions sharing the same haplotype are more likely to belong to the same genetic group (Grigorev et al. 2021), such as Eurasian accessions, but they are not necessarily from the same local population (Kuo et al. 2006). In addition, we demonstrate that linkage disequilibrium between telomeric repeat arrays and more proximal non-coding regions, previously described for single chromosome ends in humans and *A. thaliana* (Kuo et al. 2006; Baird et al. 2000; Baird et al. 1995), as a common feature at all non-rDNA chromosome ends in *A. thaliana*. The mitochondrial DNA insertion event observed in nine Eurasian accessions is a good example for summarizing these patterns in conjunction with the mutational process we propose (Supplemental Fig. S6). The ten accessions, from different localities, contain a conserved mitochondrial fragment and highly similar repeat-adjacent sequences, but the repeat arrays differ in sequence. A likely scenario is that the mitochondrial fragment was inserted before these ten chromosome ends diverged (Kuo et al. 2006). Base substitutions in the telomeric repeat arrays then occurred stochastically in different accessions during repeat amplification.

Our analysis of fixed differences in very closely related individuals has shown that telomeric repeats experience apparently much higher mutation rates than high-complexity sequences in chromosome arms, especially when it comes to repeat number. Telomeric repeats are therefore potentially helpful when attempting to reconstruct relationships between closely related individuals at high resolution. Information from telomeric repeats might become particularly useful if combined with genome-wide analyses of microsatellite and minisatellite mutations (Marriage et al. 2009). The substantial intra-individual variation in telomeric repeats also offers opportunities for studying the mechanisms of replication slippage and unequal crossing over of minisatellites (Smith 1976), given that the entire telomeric repeat arrays can be confidently captured by single HiFi reads.

In addition to the hypervariable arrangements of degenerate and variant telomeric repeats, we discovered instances in chr2q of Dralll-1 and chr3q of Cas-6 where larger fragments that included both specific arrangements of variant repeats and unique sequence fragments were duplicated in head-to-tail fashion. These duplications could be the result of simple expansion by the same mechanisms that cause other HORs. Alternatively, they could be caused by telomerase-independent repair triggered by a double-strand break (DSB; Kim et al. 2020). At the end of chr2q of Hum-2, a unique sequence fragment interrupted the variant telomeric repeats, and a related arrangement was found at chr5q of the same accession. This observed configuration may have been generated after a DSB event by a recombination process referred to as chromosome healing (Kim et al. 2019; Baird 2018; Ballif et al. 2004). These structures are reminiscent of a few instances of extensive monomer homogenization we found, which might involve ALT as a telomerase-independent process (Geiller et al. 2022; Marzec et al. 2015).

Our study leaves several open questions for future studies. The first question is where the genuine boundary between regions maintained by telomerase and maintained by DNA polymerase lies. Since the telomeres of telomerase-deficient mutants progressively shorten only for a few generations, after which no further shortening is observed (Chaux et al. 2023), we hypothesize that a fraction of canonical repeats is not involved directly in telomere maintenance. The next two questions concern the two chromosome ends next to the 45S rDNA clusters. One challenge will be to accurately assign telomeric reads adjacent to rDNA to specific chromosome ends, which is hampered here by a lack of complete assemblies of rDNA arrays in diverse genomes (Fultz et al. 2023). Another one is why there are significantly fewer variant repeats present in these two chromosome ends compared with the other eight non-rDNA chromosome ends. Lastly, the sharing of unique sequences across chromosome ends points to mechanisms for healing of broken chromosome ends by telomere capture (Baird 2018). The sequence structures we have described in the germline facilitate the question to be investigated in detail.

## METHODS

### HiFi-based data collection

Forty-eight HiFi-based assemblies and read sets, representing 46 natural accessions, were obtained from five public sources. The datasets of 44 accessions were from Wlodzimierz et al. (2023), the Kew-1 accession was from Christenhusz et al. (2023), and three independent Col-0 datasets were from Rabanal et al. (2022), Wang et al. (2022), and Naish et al. (2021).

Three HPG1 accessions (Exposito-Alonso et al. 2018) were sequenced with one SMRT Cell on the Sequel II platform (PacBio). Plant growth (Contreras-Garrido et al. 2023), DNA extraction from a single plant (Wlodzimierz et al. 2023), preparation of a multiplexed sequencing library followed by HiFi sequencing (Rabanal et al. 2022), and genome assembly (Wlodzimierz et al. 2023), were performed as previously described.

Of the 49 wild *A. thaliana* accessions, 33 were from Eurasia and three from North America. Thirteen accessions were identified as relicts (Wlodzimierz et al. 2023), of which nine were from the Iberian Peninsula, and the remaining four were from Madeira, Cape Verde, Italy and Tanzania.

### Extraction of telomeric sequences

In *A. thaliana*, two out of ten chromosome ends have large 45S rDNA repeat arrays adjacent to the telomeric repeats, causing most assemblies collapse and thus preventing correct mapping of telomeric sequences (Fultz et al. 2023; Copenhaver and Pikaard 1996). Two alternative strategies were employed to extract telomeric sequences, depending on whether the sequence was adjacent to long 45S rDNA sequences.

For the eight non-rDNA chromosome ends, an alignment-based approach was employed. For each sample, HiFi reads were aligned to the corresponding assemblies using minimap2 v2.24 (Li 2018) with the parameter -ax map-hifi. The output SAM files were converted to BAM format using Samtools v1.10 (Li et al. 2009) functions view -Sb, sort and index. Since the repeat-adjacent regions of different chromosome ends, which serve as markers for uniquely anchoring reads, were known to be similar in sequence (Heacock et al. 2004), all-against-all pairwise alignments of the 5 kb sequence adjacent to the telomeric repeats were performed for each chromosome end with BLAST v2.13.0+ (Altschul et al. 1990). This resulted in a maximum alignment overlap of 3 kb. Therefore, only reads containing at least 4 kb of repeat-adjacent sequence were extracted with samtools view -hb -L (Farrell et al. 2022; Tan et al. 2022; Grigorev et al. 2021). BAM files were converted to FASTA format using samtools bam2fq and processed with seqtk v1.3 (https://github.com/lh3/seqtk) using option seq -A. Potentially chimeric reads and reads containing sequencing errors were excluded after visual inspection. All reads were manually clipped to remove non-repeat sequences, retaining only the telomeric tracts. Since the irregular degenerate repeat content made the boundary between the non-repeat and repeat portions ambiguous, the start of the telomeric repeat array was arbitrarily defined as the first instance of the sequence (T)_x_(M)(G)_y_(M) (M = A or C).

For the chr2p and chr4p ends, which contain large 45S rDNA arrays, reads were directly extracted without help of the corresponding genome assembly. Using minimap2, reads that aligned to the 45S rDNA sequence of Col-0 (Rabanal et al. 2022) were identified. Reads with at least three consecutive telomeric repeats were further retained. The 45S rDNA portions of these retained reads were aligned pairwise using BLAST. It resulted in the length of identical 45S rDNA sequences being either less than 4,800 bp or nearly the entire length of the query sequence. Reads with at least 5 kb of 45S rDNA sequences were thus extracted and clustered into two groups, putatively from chr2p and chr4p, per accession based on sequence similarity. Based on a 45S rDNA reference sequence (Rabanal et al. 2022), RepeatMasker v4.0.9 (Smit et al. 2013-2015) was used to mask and exclude the rDNA regions from the reads, leaving only the telomeric repeats for further analysis.

To facilitate downstream analysis, reads with telomeric repeats in the 3’-CCCTAAA-5’ orientation were first reversed to 5’-TTTAGGG-3’ using seqtk with function seq -r, followed by processing with Tandem repeats finder v4.09.1 (Benson, 1999) to identify repeat units. After manual curation, units were arbitrarily defined as beginning from the first T and ending with the last non-T base along the sequence. For example, the sequence TGTTTAGGGTCTGATGGG was split into the units TG TTTAGGG TCTGA TGGG.

### Evaluation of short homopolymer errors

Because at each end of the reads, small indels, particularly 1-bp deletions, often dominated the canonical TTTAGGG repeat regions, specifically TTAGGG (with two instead of three Ts) or TTTAGG (with two instead of three Gs), and these occurred at random positions. To determine whether these indels were caused by somatic mutations or sequencing errors (Lal et al. 2021), the correlation between the likelihood of errors and the occurrence of indels for each read was examined.

The likelihood of error was quantified based upon subread coverage and quality value of the HiFi reads. Samtools view -X followed by an awk command was used to extract the values of two tags, “np” (number of subreads) and “rq” (read quality), per read from the BAM files. To calculate the occurrence of indels, sequences were extracted from the read end until the variant repeat preceding the canonical repeat array. The length of each extracted sequence was divided by seven to obtain an approximate repeat number, and a hill-climbing algorithm was used to find the nearest integer that represented the canonical repeat number in the ideal sequence, minimizing the Levenshtein distance (L-distance) between the extracted sequence (the observed string) and the ideal sequence consisting entirely of canonical repeats (the expected string), obtained with the R package stringdist (Supplemental Fig. S10). This minimized distance was further divided by the length of the extracted sequence to determine the relative distance as an indication of indel density. The correlation between the likelihood of errors and the occurrence of indels for each read was plotted using R package ggplot2, and the R function cor was used to calculate the Pearson correlation coefficient.

### Identification of telomeric repeat content

To visualize the degenerate and variant repeat arrays for each accession and each end, consensus sequences were generated. Conserved units between reads were retained, while nonconserved units were marked as “N”. The frequency of occurrence for each unit type was subsequently calculated. The positions of the top 15 enriched unit types were then emphasized with different colors.

In addition, non-repeat sequences that disturbed the repeat arrays were manually extracted. Using BLAST, the sources of these non-repeat sequences were determined with TAIR10 transposon and organellar DNA sequences (Lamesch et al. 2012), as well as a library of *A. thaliana* rDNA and centromere sequences (Rabanal et al. 2022).

### Identification of telomerase RNA template sequence

In *A. thaliana*, the addition of telomeric repeats is directed by a 9-bases template 3’-UCCCAAAUC-5’ in the telomerase RNA, corresponding to 3’-TCCCAAATC-5’ in the genome (Song et al. 2019). To investigate whether the variants we observed were caused by mutations in the template sequence, all 49 assemblies were searched using BLAST with the sequence of the telomerase RNA locus of Col-0 as a query (Fajkus et al. 2019). Corresponding sequences were extracted using bedtools v2.27.1 (Quinlan and Hall 2010) with function getfasta and used as input for a multiple sequence alignment with Clustal Omega (Sievers et al. 2011). The sequence logo was visualized with WebLogo (Crooks et al. 2004).

### Estimation of telomeric repeat variants in HPG1 accessions

Three HPG1 accessions were sequenced. To assess the repeat number variation, the length of the sequences containing degenerate and variant repeats was calculated for each read with an awk script. The significance of the difference in length between accessions was evaluated with a F-test using the R function var.test. The length of each read was plotted using ggplot2.

### Estimation of telomeric repeat variants in different Col-0 datasets

Three datasets of the Col-0 accession (Rabanal et al. 2022; Wang et al. 2022; Naish et al. 2021) were compared using the methods described above. The R function var.test was additionally used to assess whether different sequencing strategies (single-plant versus pooled) affected the distribution of repeat number variation of HiFi reads.

### Haplotype structure analysis of the repeat arrays and their adjacent non-coding regions

For telomeric repeat arrays, a repeat compression approach for each sequence was used (Rautiainen et al. 2023), in order to reduce the complexity arising from repeat number variation. Pairwise L-distances between compressed arrays were calculated to estimate their similarity, followed by an F-test to assess whether there were significant differences in the similarity levels when comparing the same and different chromosome ends, and comparing the same and different genetic groups.

To identify the more proximal non-coding regions, Liftoff v1.6.3 (Shumate and Salzberg 2021) was used in conjunction with the TAIR10 gene set (Lamesch et al. 2012) to annotate the most terminal gene in the eight non-rDNA chromosome ends (Vrbsky et al. 2010). Subsequently, the fragment between the most terminal gene and the first telomeric repeat was extracted using bedtools getfasta. Multiple sequence alignment and NJ clustering of non-coding sequences was performed for each end with Clustal Omega, and pairwise L-distances were calculated using R package stringdist.

To determine whether there was any correlation between variation in the telomeric repeat arrays and the non-coding regions, the pairwise L-distance values for both the repeat and the non-coding regions were merged into a square matrix. The order of accessions for each chromosome end was determined based on the NJ clustering of the non-coding regions. Pairwise L-distances below 50 were highlighted in the matrix.

## ACKNOWLEDGEMENTS

We thank Alejandra Duque, Zhigui Bao, Katerina Romanova, Adrian Contreras, Patience Chatukuta for comments on the manuscript, and Haim Ashkenazy for discussion. This work was supported by the Max Planck Society (DW).

## AUTHOR CONTRIBUTIONS

YT designed the project. AM, CL, YT and GS performed plant growth, HMW DNA extraction and library preparation for three HPG1 accessions. FR and YT performed genome assembly. YT and WX conducted data analysis. DW provided general advice. YT and DW interpreted the data and wrote the manuscript with input from all authors.

## DATA AVAILABILITY

The datasets of three HPG1 accessions will be made available upon publication. All other datasets are publicly available.

## COMPETING INTERESTS

DW holds equity in Computomics, which advises plant breeders. DW also consults for KWS SE, a plant breeder and seed producer with activities throughout the world. All other authors declare no conflicts.

