## Supplemental Figures for "Atlas of telomeric repeat diversity in *Arabidopsis thaliana*"

### SUPPLEMENTARY FIGURES

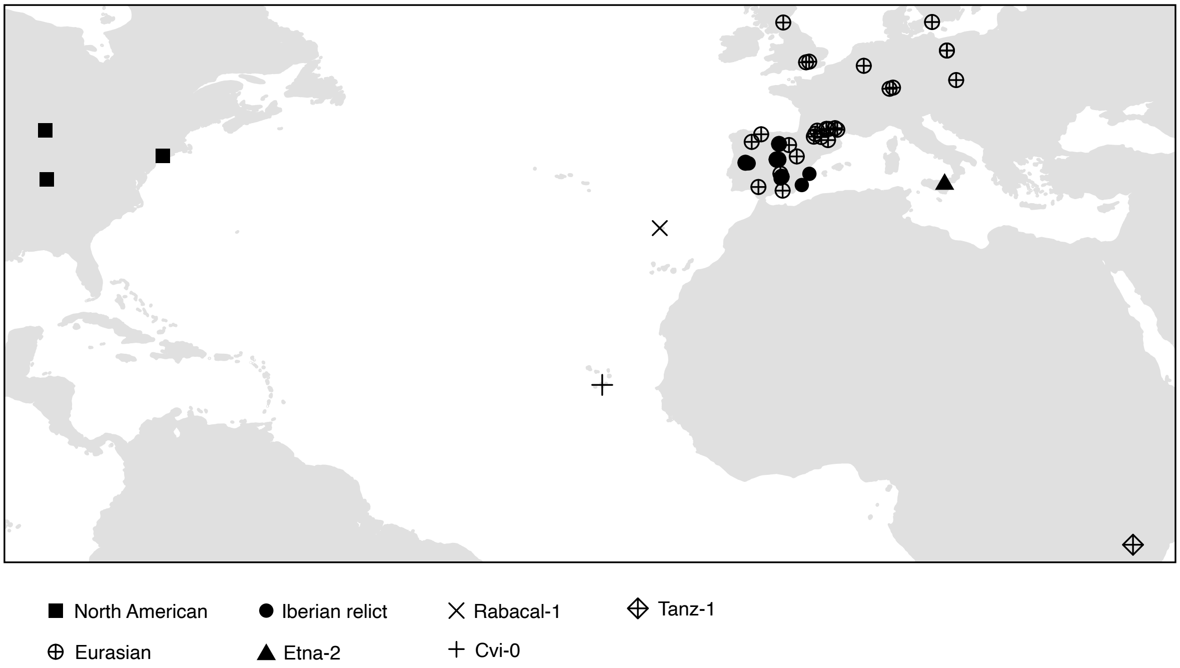


**Figure S1.** Geographic distribution of the 49 *A. thaliana* accessions analyzed. The attribution to major genetic groups was inferred by Wlodzimierz et al. (2023). Four individual accessions are directly labeled with their names.

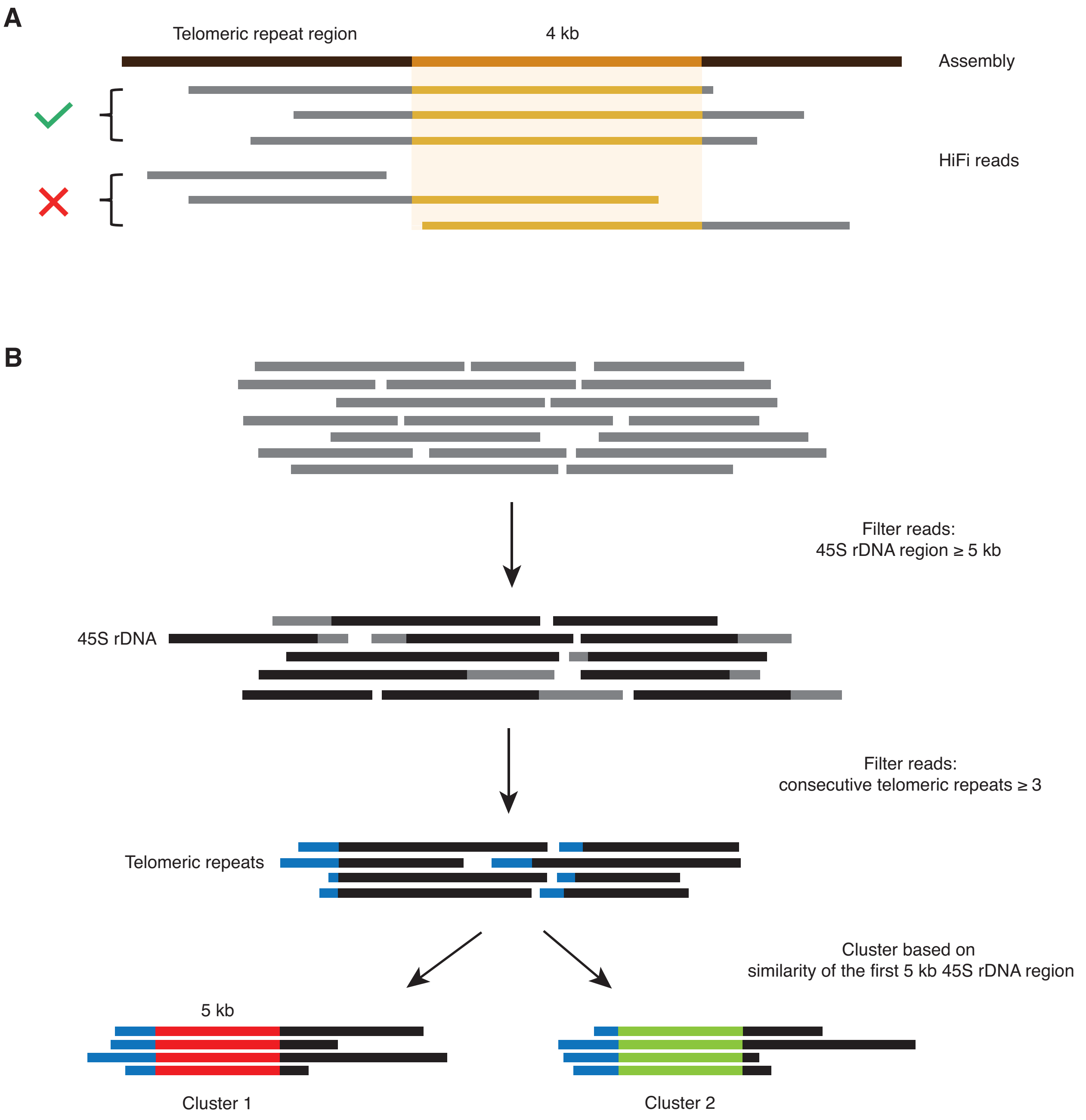


**Figure S2.** Schematic illustration of the strategies for extracting telomeric reads. **(A)** Strategy for the eight non-rDNA chromosome ends. Telomeric reads that contained at least 4-kb repeat-adjacent sequences from the relevant genome assembly were extracted. **(B)** Strategy for the two rDNA-binding chromosome ends. Reads containing 45S rDNA sequence and more than three consecutive telomeric repeats were extracted without the help of genome assemblies and clustered into two groups based on sequence similarity of the 45S rDNA region.

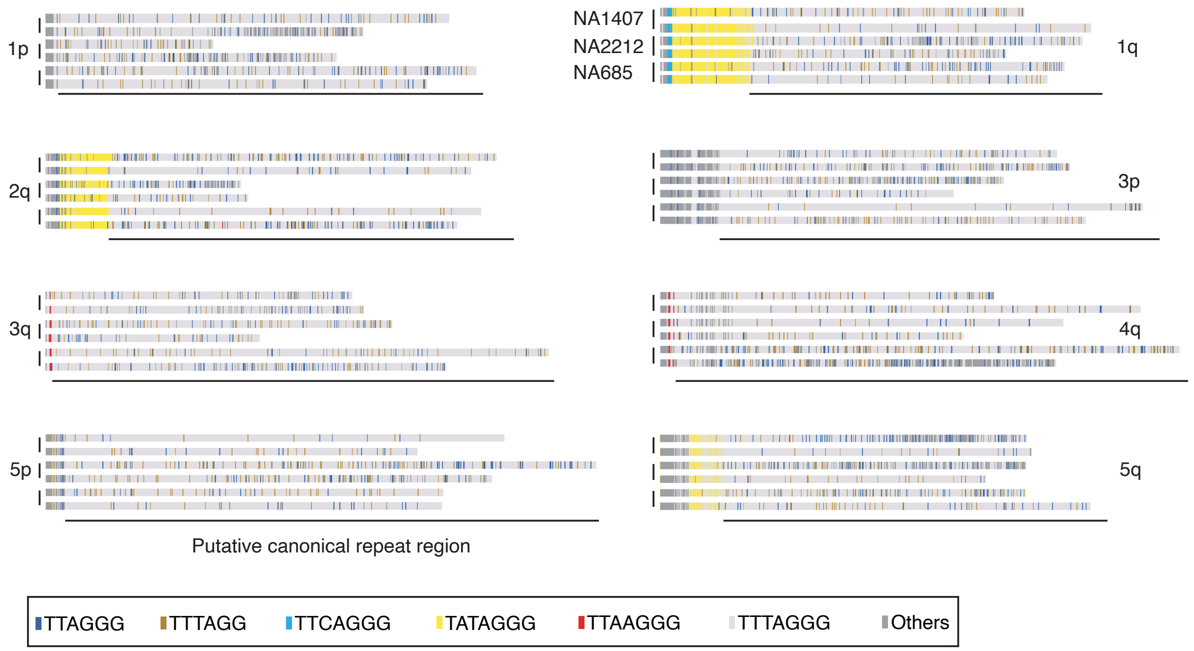


**Figure S3.** Sequence tracks showing the entire telomeric repeat arrays in the eight non-rDNA chromosome ends of the three North American accessions from degenerate, variant to canonical repeats (from left to right). Two reads were randomly selected per accession per end. Black lines at the bottom indicate the putative canonical repeat regions.

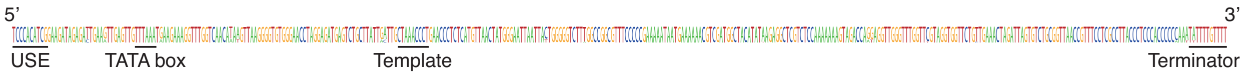


**Figure S4.** Sequence logo of the telomerase RNA locus of the 49 accessions. The conserved 9-bp template region is indicated.

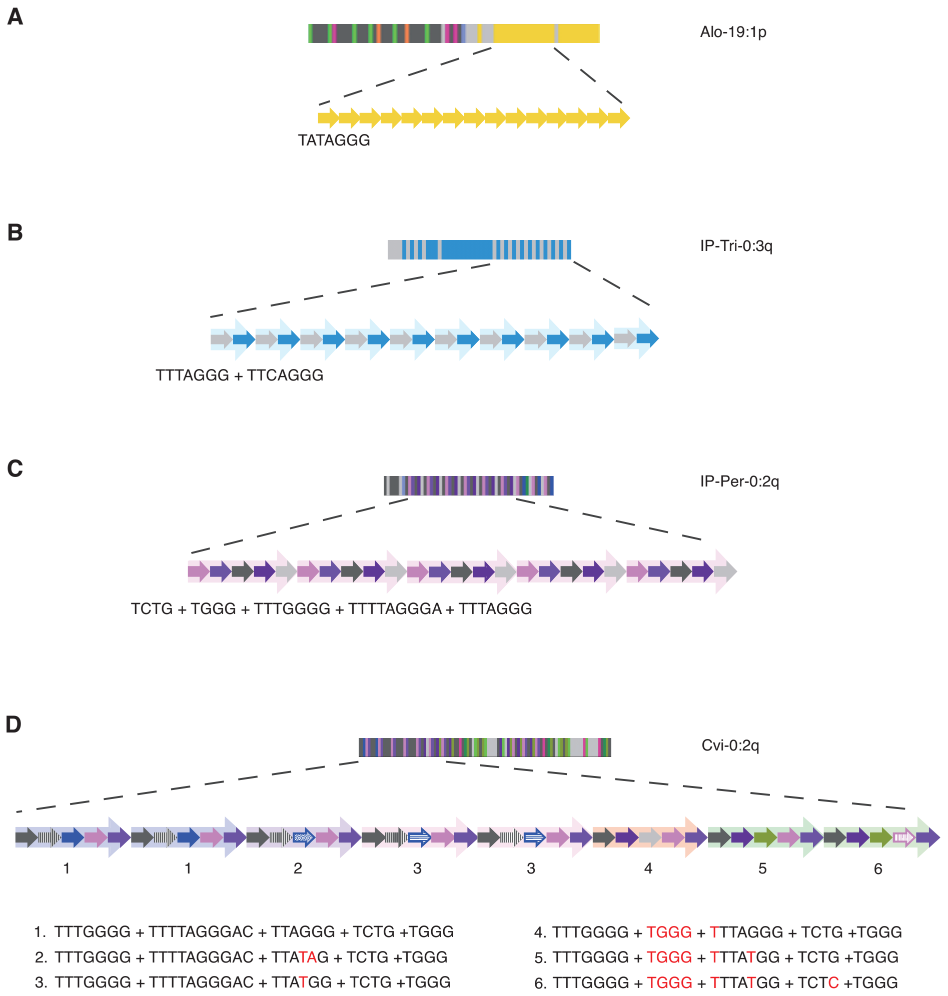


**Figure S5.** Close-up of four major types of sequence organization in the telomeric repeat arrays. Color code corresponds to that of Figure 2. **(A)** An example of monomer homogenization. In this case, a single unit is repeated 15 times. **(B)** An example of simple block expansion. In this case, two units form a block and the block is repeated ten times. **(C)** An example of expansion of identical higher-order repeats (HORs). A HOR consisting of five distinct units is repeated five times. **(D)** An example of HORs with small sequence differences. Numbers indicate different HORs.

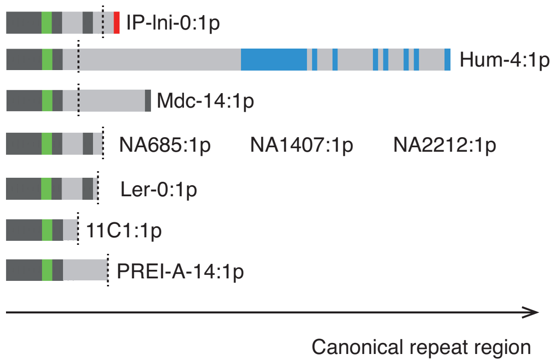


**Figure S6.** Illustration of a mitochondrial DNA insertion in nine accessions. The degenerate and variant repeat arrays are as shown in Figure 2. Vertical dashed lines denote the positions of the mitochondrial DNA fragments.

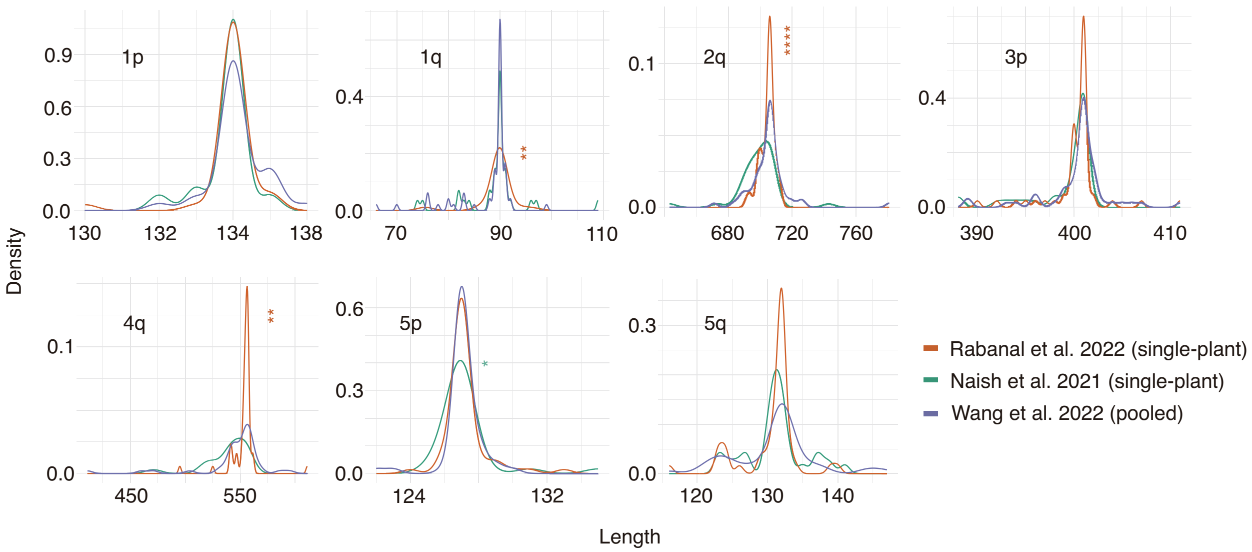


**Figure S7.** Density plot of the length distribution of degenerate and variant repeat regions at seven non-rDNA chromosome ends in three Col-0 datasets. Statistically significant differences in the deviation degree are indicated (****P<0.00001, ***P<0.0001, **P<0.001, *P<0.01).

TTCAGGGTTCAGGGTTCAGGGTATAGGGTATAGGGTTTAGGGTTTAGGGTATAGGGTATAGGGTTTAGGG

↓

TTCAGGGTATAGGGTTTAGGGTATAGGGTTTAGGG

↓

TTCAGGGTATAGGGTTTAGGG

**Figure S8.** Schematic representation of the repeat compression process.

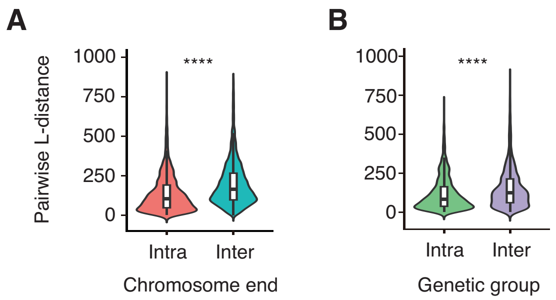


**Figure S9.** Violin plots showing the distribution of pairwise L-distances. Statistically significant differences between accessions are indicated (****P<0.00001). **(A)** Comparison of values within and between chromosome ends. **(B)** Comparison of values within and between genetic groups.

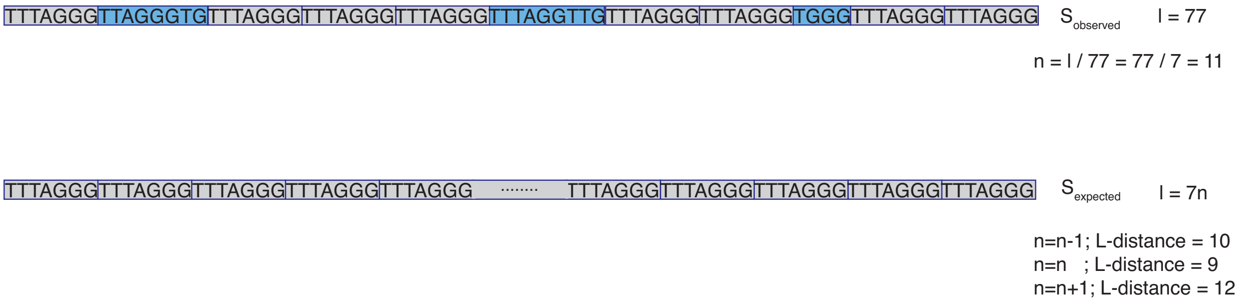


**Figure S10.** Example of the process for determining the expected string and calculating the L-distance, which represents the occurrence of indels. Blue blocks indicate variant repeats, while gray blocks indicate canonical repeats. ‘S’ indicates ‘string’, ‘n’ should be an integer.
